# pISA-tree - a data management framework for life science research projects using a standardised directory tree

**DOI:** 10.1101/2021.11.18.468977

**Authors:** Marko Petek, Maja Zagorščak, Andrej Blejec, Živa Ramšak, Anna Coll, Špela Baebler, Kristina Gruden

## Abstract

We developed pISA-tree, a straightforward and flexible data management solution for organisation of life science project-associated research data and metadata. pISA-tree was initiated by end-user requirements thus its strong points are practicality and low maintenance cost. It enables on-the-fly creation of enriched directory tree structure (<u>p</u>roject/<u>I</u>nvestigation/<u>S</u>tudy/<u>A</u>ssay) based on the ISA model, in a standardised manner via consecutive batch files. Templates-based metadata is generated in parallel at each level enabling guided submission of experiment metadata. pISA-tree is complemented by two R packages, *pisar* and *seekr*. *pisar* facilitates integration of pISA-tree datasets into bioinformatic pipelines and generation of ISA-Tab exports. *seekr* enables synchronisation with the FAIRDOMHub repository. Applicability of pISA-tree was demonstrated in several national and international multi-partner projects. The system thus supports findable, accessible, interoperable and reusable (FAIR) research and is in accordance with the Open Science initiative. Source code and documentation of pISA-tree are available at https://github.com/NIB-SI/pISA-tree.

## Introduction

Compared to a decade ago, present-day life science projects generate vast amounts of data, that are frequently heterogenous. Both researchers and funders are aware that published results often lack important information that would ensure their reproducibility. Raw data supporting researchers’ conclusions are frequently either inaccessible, inadequately organised or not sufficiently annotated to allow straightforward reuse^1–3^. To alleviate these issues, the FAIR data principles^4^ that advocate policies which ensure findability, accessibility, interoperability and reusability of the generated data were proposed. Thus, reporting, format and semantic standards are of utmost importance^5^. Minimum information metadata standards, providing guidelines for standardised reporting of experimental results, have been developed for a large number of wet-lab assays; e.g. plant phenotyping (MIAPPE^6^), quantitative PCR (MIQE^7^), gene expression microarrays (MIAME^8^), proteomics (MIAPE^9^); as well as dry-lab assays, e.g. quantitative modelling (MIRIAM^10^) and simulations in systems biology (MIASE^11^). To organise and keep track of experimental metadata, the ISA framework^12^ implemented the ISA model that captures experimental metadata on three hierarchical levels: Investigation, Study and Assay (abbreviated as ISA levels). Each ISA level contains files describing experimental goals, ontologies used, experimental design, protocols, and experimental conditions. ISA framework developers also provide ISA tools^13^, a software suite that, among others, contains a metadata editor and packages for integration with common programming languages^14–16^. ISA tools use a series of graphical interface input forms to generate metadata in the ISA-Tab or ISA-JSON format and use assay-specific templates covering a wide variety of methodologies and community standards. In parallel, based on minimum information standards, several public repositories^17^ have formulated metadata templates, that guide the researchers before submitting the data with publications. Besides methodology-specific repositories, there are general repositories that offer long-term research data and metadata storage following FAIR principles. One such service is Zenodo (www.zenodo.org), an open science repository developed by CERN that accepts datasets in any format. Another is SEEK/FAIRDOMHub^18^, a general repository that adopted the ISA levels to structure project-related data. While these all fullfill the requirements of OpenScience they are difficult to embrace and adopt by the researchers that are involved in data acquisition.

For the duration of a research project, datasets are frequently stored and analysed locally, usually on the researcher’s personal computer or institution’s network drive. Institutions have various strategies on how to deal with organisation and medium to long-term storage of project data. These strategies may include the use of a Laboratory Information Management System (LIMS), electronic laboratory notebook (ELN), and either centralised or decentralised storage of raw and analysed data. However, for research projects, such software solutions can have some disadvantages -they might be associated with license costs, are prone to rigidity, can become unsupported over the course of a few years, or are unable to handle diverse and large scale datasets^19,20^. Several services were also developed for FAIR data managmenet, besides SEEK/FAIRDOMHub^18^, COPO^21^ and e!DAL^22^ are examples of such services. In principle, those services can also be run locally at an institutional level; however, this requires substantial IT support and discipline in data upload. Project managers are thus more likely to use these services only for data deposition for long-term storage at the end of the project than for immediate data storage.

To address all the issues mentioned above, we have developed pISA-tree, an easy-to-use and maintain system for intra-institutional organization and structured storage of research data, with a special emphasis on the generation of adequate metadata. pISA-tree uses the ISA levels, FAIR principles, and minimum information standards for metadata formulation, and encourages the inclusion of metadata on-the-fly, when the experiments are designed. Its complementary R packages enable easy transfer to public (meta)data repositories and (re)use of data in analytical pipelines.

## Results

We developed pISA-tree and its accompanying R packages (*pisar* and *seekr*) to provide a user-friendly system that guides researchers towards organisation of their data without the requirement of any advanced systems maintenance. It was designed by data scientists in constant interaction with reserchers involved in data acquisition to allow for adoption by both.

### pISA-tree functionalities

pISA-tree is based on a consecutive set of batch files that generate an enriched directory tree structure for scientific research projects. It uses a familiar directory system with a standardised structure to organise each **<u>p</u>**roject hierarchically into three nested ISA levels: **<u>I</u>**nvestigations, which consist of multiple **<u>S</u>**tudies, where each individual study can contain multiple **<u>A</u>**ssays conducted on a set of samples from a single coherent experimental setup (Fig. 1, Supplementary File 1). Typically, an investigation aligns well with a defined experimental question or project work package, while a study encompasses a sample collection from one experiment.

**Fig. 1.**
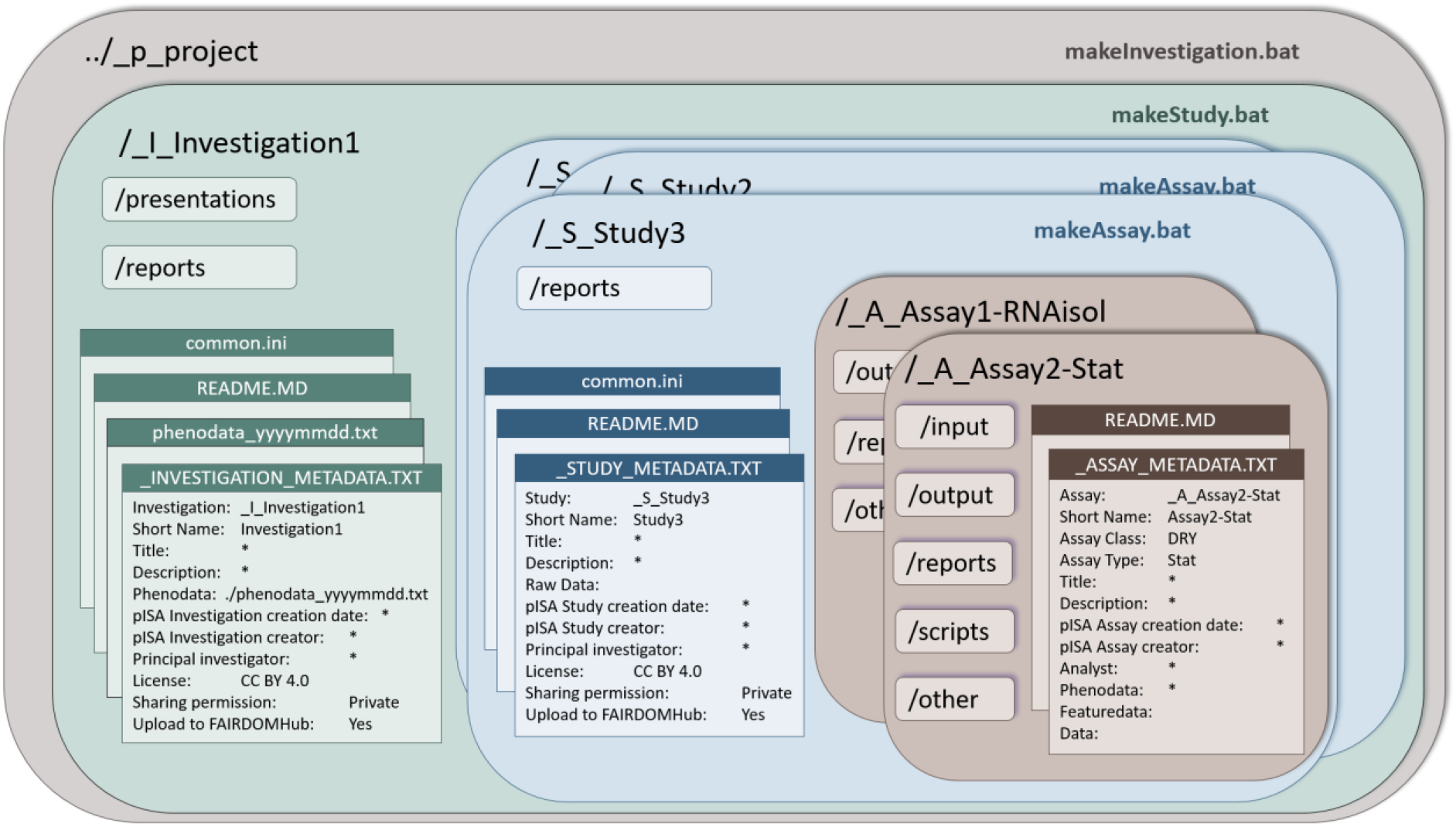
pISA-tree directory tree structure and corresponding metadata files. Large dark-shaded rectangles with rounded corners represent pISA-tree levels and corresponding directories, light-shaded rectangles with rounded corners represent supplementary subdirectories as predefined for each level, and rectangles represent pISA-tree related files. Bolded file names in the upper right corner of the levels represent batch files (.bat) that generate new pISA-tree levels.

One of the core components of the system are consistent metadata files, associated with each pISA-tree level (Fig. 1). Prior to folder creation, the user answers a series of questions (keys) with predefined attributes or free-text answers (values), which are then stored in metadata files as tab-delimited key-value pairs. Key-value pairs can follow any controlled vocabulary^16^ and can be further edited using any text editor. This encourages users to enter the minimal information at an early stage of research preparation and provides the advantage of finding relevant information later. In addition, updated and consistent metadata can be used in the data analysis phase.

While the upper levels, i.e. project, investigation and study, hold a more descriptive role, the assay level is the richest level of the research record. Each data analysis step can be defined as a separate Assay. Moreover, pISA-tree can support two different classes of assays: laboratory experiment (wet) and data analysis (dry). Each assay class contains several pre-prepared assay types, depending on measurement methodology (e.g. RNA isolation, qPCR, GC-MS). These templates were developed by taking into account minimal information standards (e.g. MIQE^7^ for qPCR) as well as consulting researchers performing the assays. The assay class and type define the subdirectories and other standardised information that is typically required. For example, besides common subdirectories such as `output` and `reports`, the wet-lab assays will have the `output\raw` subdirectory for storing raw experimental data, whereas the dry-lab assays will have the `scripts` subdirectory (Table 1). All options and predefined minimal information are kept in text files and can be customised to extend assay classes and types. For each new class or type, the user adjusts the directory structure and defines metadata template files.

**Table 1.**
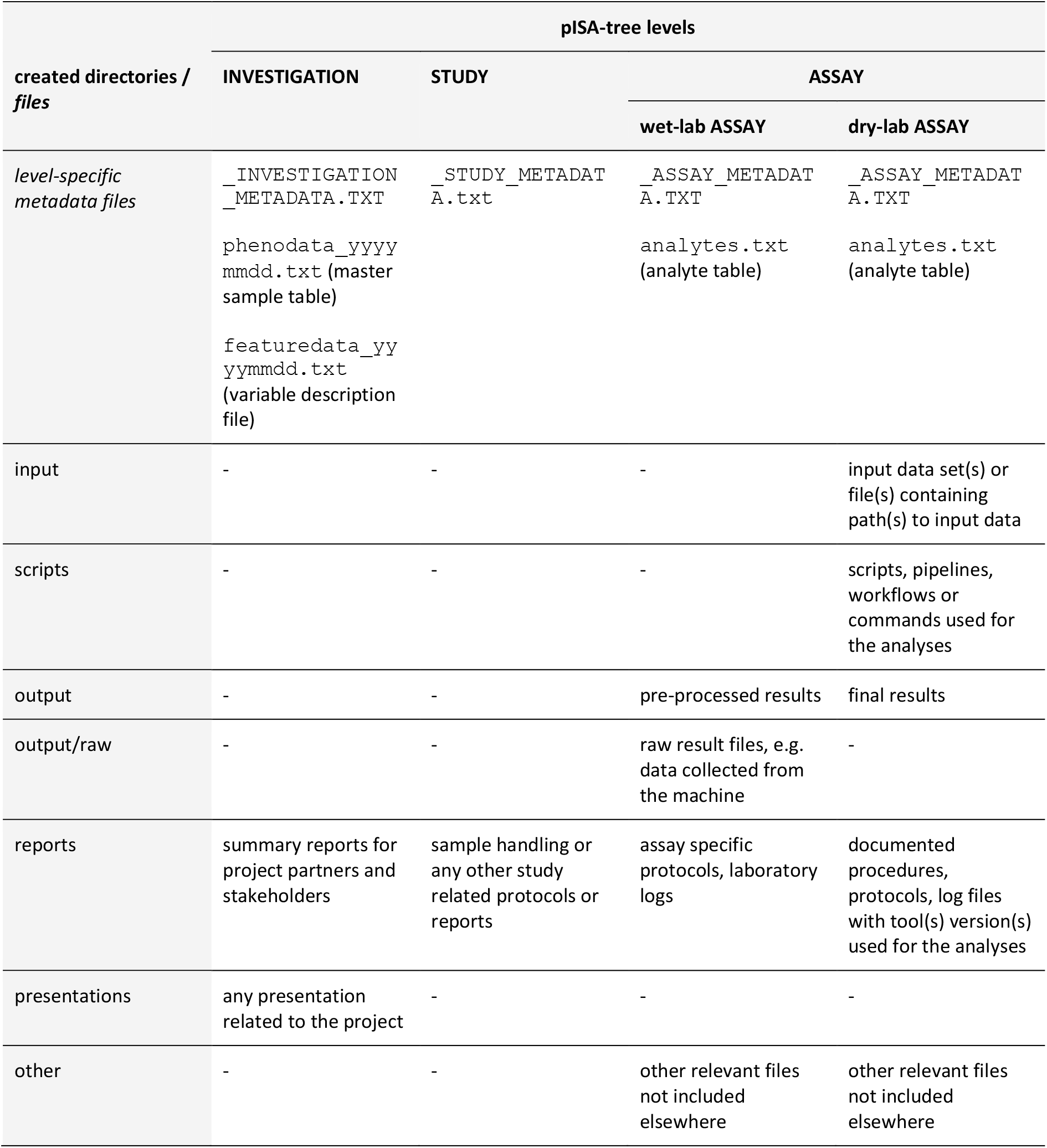
Automatically generated pISA-tree directories and metadata files and recommended allocation of data files. Each level contains level-specific metadata files and directories containing data. ‘-’ directory not created at this level.

### Additional pISA-tree features for metadata management

#### Common metadata

Before creating any project subdirectories, key-value pairs that should be present in all sublevel metadata files, e.g. organisation or license information, can be added to the `common.ini` file. This way, key-value pairs will be automatically assigned to all directories within the project and do not need to be re-entered for each new investigation or study.

#### Samples description

The investigation level specifically holds the master sample description table, namely the phenodata file (`phenodata_yyyymmdd.txt`), which should be populated with all relevant information on all the experimental samples that will be collected in the associated studies and further analysed in corresponding assays. The phenodata file should therefore contain the experimental design information required for planned data analysis. We advise users to take the requirements of the community-specific minimal information standards into account when preparing the phenodata files, i.e. use MIAPPE^5^ for plant samples. More details on how to structure and keep track of the potential changes in the phenodata file can be found in the pISA-tree user manual (Supplementary File 1).

#### Experiment description

Users are also encouraged to create descriptions of all measured variables within assays and store them in a featuredata file (`featuredata_yyyymmdd.txt`). Certain assay types additionally generate an analyte table (`analytes.txt`) which serves as a template for input of specific measurements (e.g. sample quality control values), thus helping the wet-lab researchers to input the results of experiments into the system.

#### Consistency and overview

The metadata files created are machine-readable, enabling interoperability and interconnectedness across research fields. All levels contain three auxiliary batch scripts: `showtree.bat` displays the complete directory tree structure and allows for a better overview of the information in the system for complex projects; `showmetadata.bat` displays all recorded metadata thus far; and `xcheckMetadata.bat` displays missing mandatory metadata fields. These auxiliaries thus help the scientists when reviewing their project status in terms of data management completeness, thus promoting data FAIRness.

### Integration of pISA-tree into web-based data management systems

Because the R environment for statistical computing has a firmly established developer and user community for bioinformatic data analysis, with well-maintained packages, we further complemented pISA-tree with two R packages. The predefined pISA-tree directory structure enables integration into automated data analysis pipelines, while R enables repeatability of analyses, e.g. by including output of the *SessionInfo()* function in the analysis reports or using R project environment. Therefore, we further complemented pISA-tree with *pisar*, an R package that enables import of pISA-tree metadata into R workflows (Fig. 2) and thus repeatability of R analyses within the pISA-tree system. Since information is structured in a standardised way and augmented by metadata, connection to other local or cloud-based FAIR data management platforms can be established. The other part of the system enables seamless upload to the project metadata repository FAIRDOMHub^18^ using the R package *seekr* (https://github.com/NIB-SI/seekr) which interacts with the FAIRDOMHub API. When using general data repositories for long term storage, *pisar* package allows conversion of pISA-tree metadata into ISA-Tab format, which we recommend to upload together with an archived file containing the entire project (Fig. 2).

**Fig. 2.**
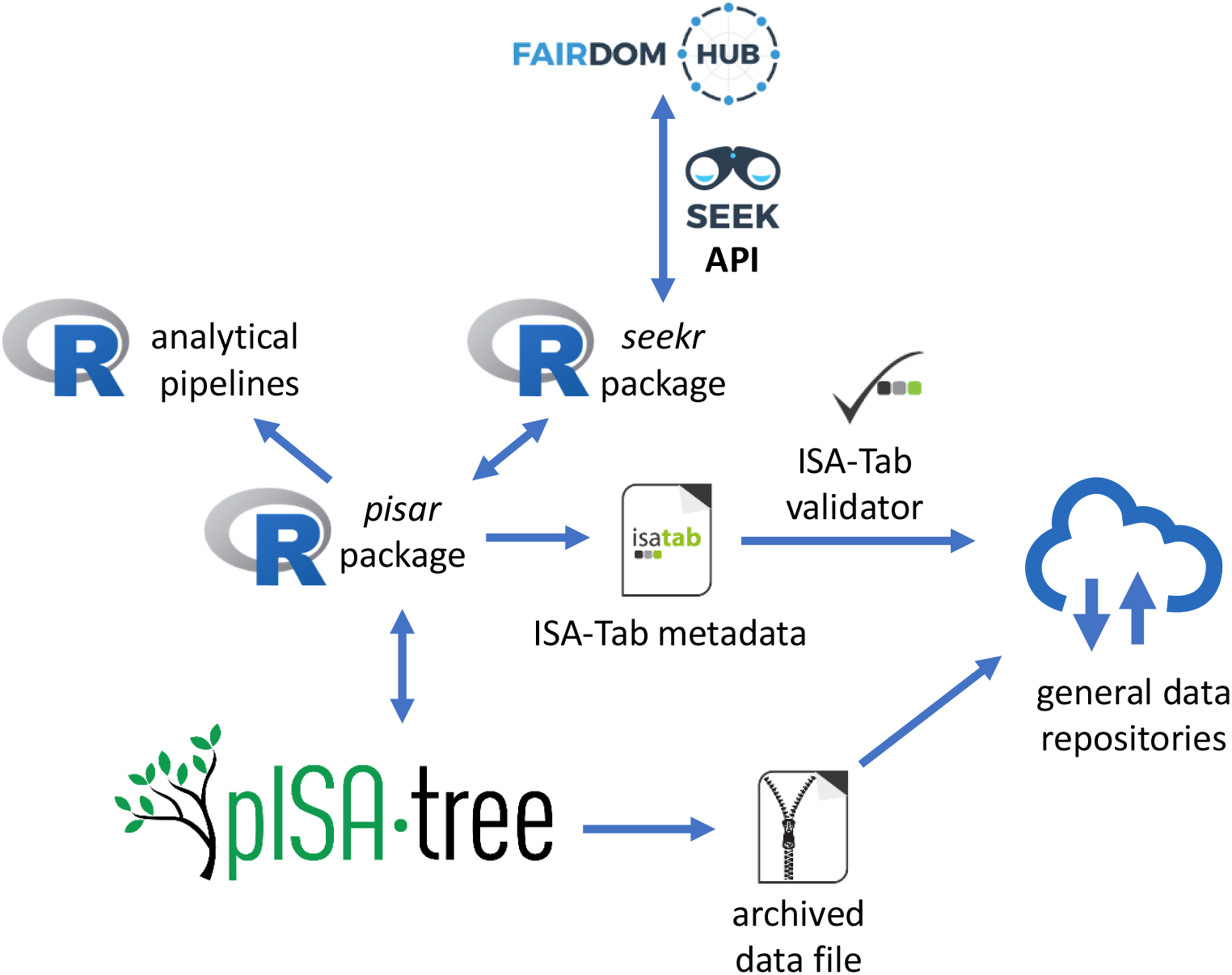
pISA-tree information flow between public metadata repositories and R analytical pipelines. Data flow between pISA-tree, R environment and the metadata repositories is represented with blue arrows. Users can acquire pISA-tree metadata for analysis in R using the *pisar* package. In addition, *pisar* enables the conversion of metadata into ISA-Tab format to validate using ISA-Tab validator and store together with the archived project data file in public repositories.

### Examples of projects managed using pISA-tree

FAIR paradigms are becoming integral to the way we conduct scientific research. We developed pISA-tree for local project data management to facilitate implementation of these paradigms in day-to-day research. We illustrate the usage of pISA-tree with two example life science projects described below, both including diverse assay types. The *seekr* package was used for the automatic upload of the complete projects to FAIRDOMHub, thus making these projects data easily accessible and reusable.

The first example is a multidisciplinary project on insect pesticide development. The project (“Using RNAi and systems biology approaches for validation of insecticide targets in CPB guts”, https://fairdomhub.org/projects/252) has both wet and dry lab assays. The scope of this project was to develop double-stranded RNA (dsRNA) based insecticide against Colorado potato beetle pest. It included identification of Colorado potato beetle gene targets, establishing dsRNA production, and validating the dsRNAs’ insecticidal potential in laboratory and field trials^23^. The directory tree of this project consists of three investigations covering altogether 16 studies with wet-lab and dry-lab assays (Fig. 3). The investigation “_I_01_LabTrials” includes 11 studies, the first three of which contain data on dsRNA design and production, while the rest contain data on feeding trials. The phenodata file in this investigation describes all samples in the underlying assays in detail. The “RNAisol” wet-lab type assays in the feeding trial studies include analyte files that contain information on isolated RNA quantity and quality parameters measured for samples analysed in the particular trial.

**Fig. 3.**
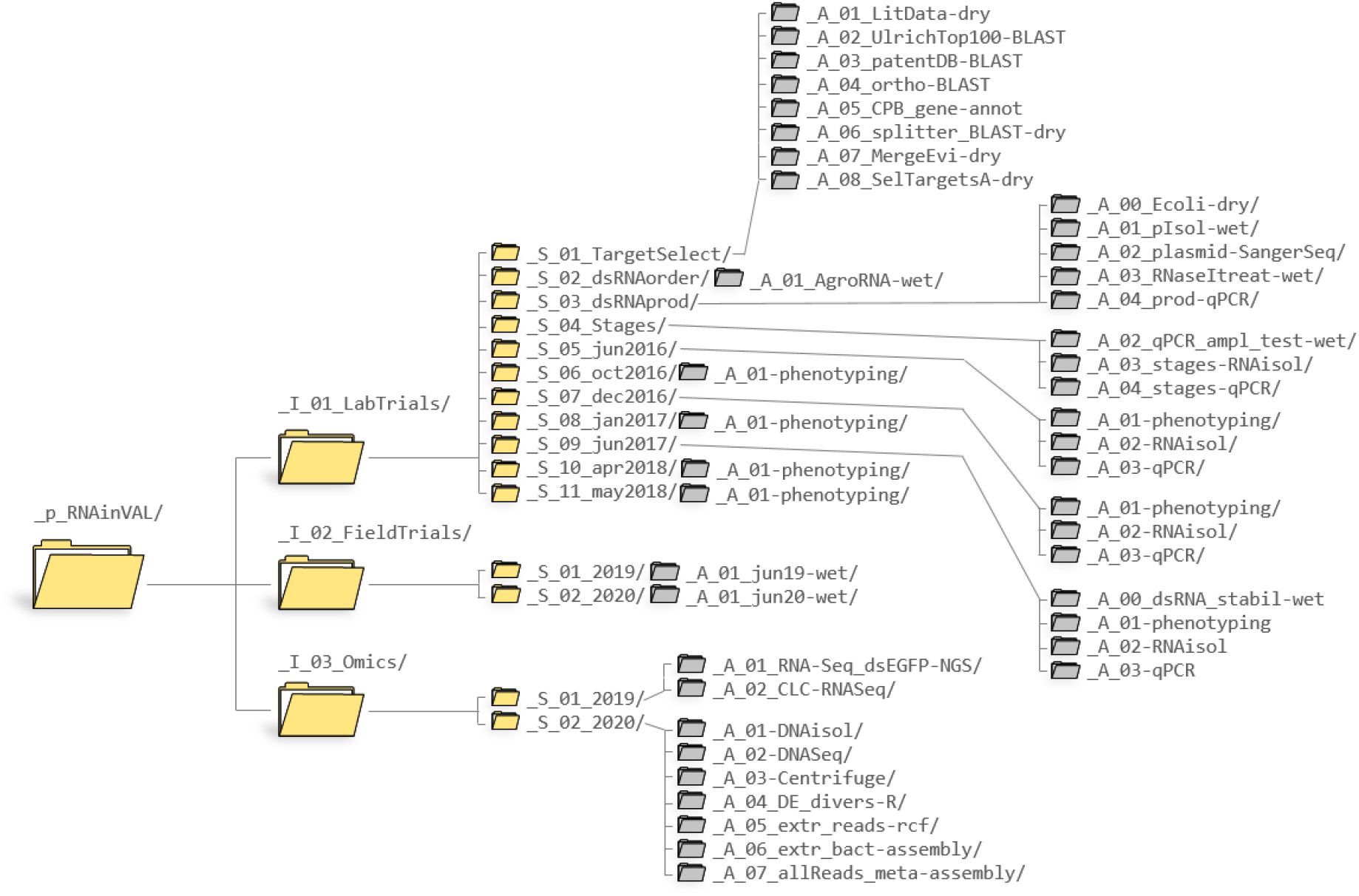
pISA-tree directory structure of the project from Example 1. Only the pISA-tree level directories are shown for clarity; with project, investigation and study directory icons in yellow and assay level icons in grey.

The second example is an implementation of pISA-tree in a large-scale dry-lab project on *de novo* potato transcriptome assembly (“*Solanum tuberosum* Reference Transcriptomes”, https://fairdomhub.org/projects/161). This project’s aim was to construct a pan-transcriptome from high-volume data accessible from publicly available resources and in-house defined workflows in a collaborative way^24^. The project consists of a single investigation, four studies and a plethora of data files (Fig. 4), encompassing four consecutive bioinformatics procedures: (1) high-throughput sequencing acquisition and pre-processing, (2) *de novo* assembly, (3) cultivar-specific transcriptomes generation and annotation, and (4) pan-transcriptome generation and annotation. Assays nested within the Studies are structured as predefined for the dry-lab assay class (see Table 1), yet no additional specific assay types were developed within this project, as they were custom for each step of the developed assembly and quality control pipeline.

**Fig. 4.**
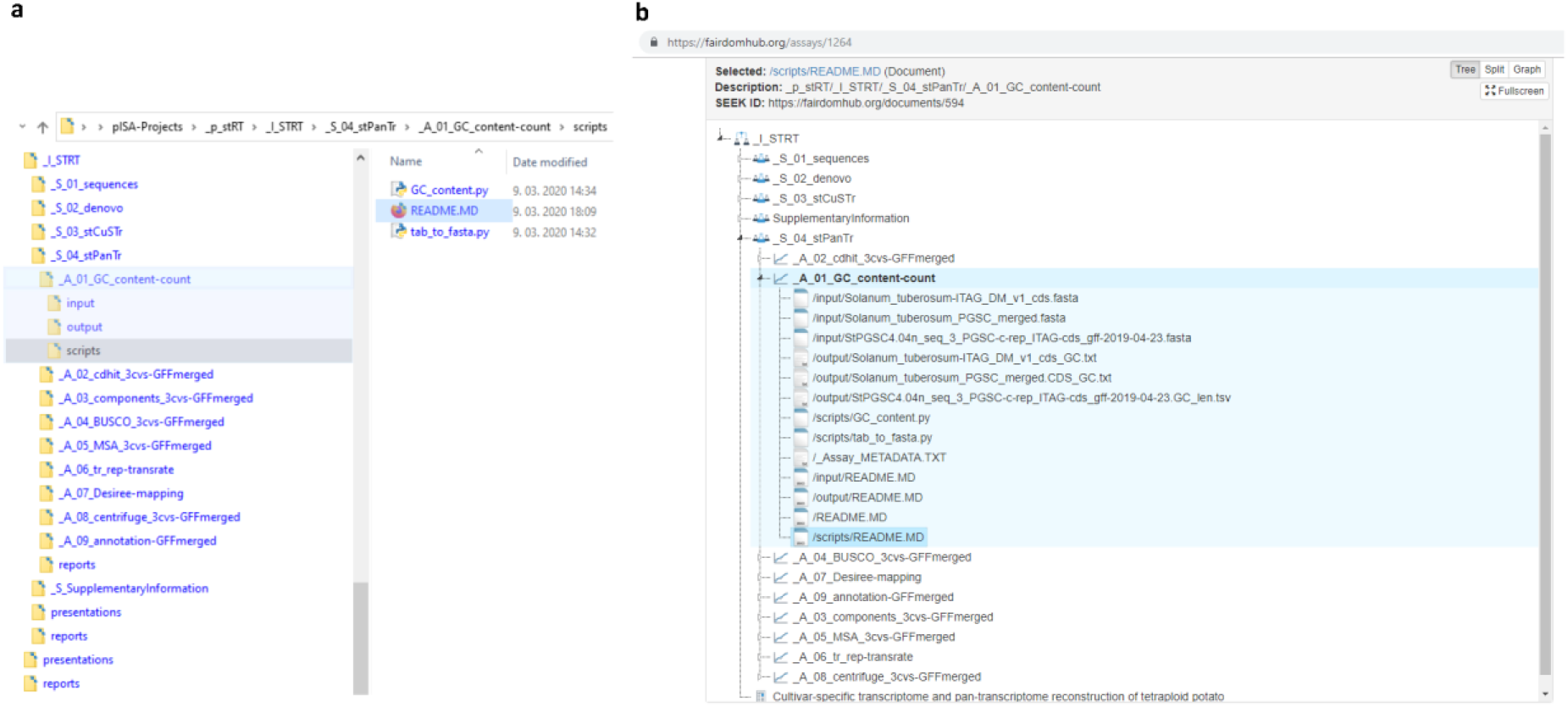
Local pISA-tree directory structure and the corresponding FAIRDOMHub structure of the investigation from Example 2. **a)** Hierarchical structure of investigation “_I_STRT” from the project of Example 2 as stored on the local network drive, with one of the assays shown in detail. **b)** Same pISA-tree structure as shown in panel a stored at FAIRDOMHub and an example of all file types within one of the assays shown in detail. Slash characters (/) in file names are used to denote the subdirectory paths of the listed files in the assay.

## Discussion

To promote the adoption of FAIR principles, many funding organisations and journals are committed to FAIR and open data principles^25,26^. Often, project proposals are expected to contain at least a draft of data management plan^27^. This plan has to incorporate policies for data collection, organisation and exchange throughout the project duration and also after its end, since it must also take into account local storage and deposition to publicly accessible repositories. The life science community has at large adopted the core FAIR values. Bottlenecks lie in the implementation of said management plans, partially because the data management systems in place are not designed for researchers that typically collect, annotate, and organise the data. As these systems are usually coordinated on an institutional level, their practicality and usefulness for individual researchers or research groups are often not taken into account^28^. In that respect, pISA-tree is distinct, initiated by end-user requirements and implemented by developers with an emphasis on practicality. The development went through several rounds of refinements to accommodate most of the involved life scientists’ needs.

Compared to other life science project data management systems, one of the main advantages of pISA-tree is its simplicity. It is based on file system directories, inherent to all operating systems and can therefore be used by scientists with basic computer skills following a short practical course or careful reading of the user manual (Supplementary File 1). Organising metadata is a fairly complex task, thus the process in pISA-tree is to some degree automated by predefined metadata keys and values. We believe that a simple system will be more easily adopted by a wide range of scientists, thus increasing compliance with FAIR principles. For the same reason, pISA-tree does not require professional technical support but we recommend a data steward being nominated to take care of maintaining order, access control, updates, uploading to public repositories, and data backup. Therefore, the data management system we propose here is particularly practical for smaller institutions or research groups with limited resources and staff. Other advantages include open source code, independence from web browsers and third-party dependencies, and straightforward access control.

pISA-tree metadata are stored in plain text files that can be edited, conferring to the system high flexibility. In addition, it is not limited to specific laboratory or computational technologies, as the provided wet and dry metadata templates can be applied and adjusted to any new technology. The user is encouraged to create the pISA-tree directory structure and corresponding metadata files before conducting the experiments. However, considering the impracticality of strict requirements for immediate metadata input, which could discourage end-users^27^, pISA-tree flexibility allows the user to complete or correct the metadata at the later stages. This is a very convenient feature trait of pISA-tree compared to other systems such as ISA tools^13^, which assume that the assays have already been carried out and the data was analysed.

Compared to ISA tools, in pISA-tree the metadata files are organised somewhat differently. ISA tools does not use a hierarchical directory structure, instead, all metadata files are stored in one directory. With larger projects hierarchical organisation of experiments helps keeping the overview and eases reuse. Another distinction from ISA tools is that in pISA-tree, sample descriptions and experimental design are stored in the phenodata file and the measured variables are stored in featuredata file at the investigation level. The structure of these files are comprehensible for wet-lab researchers. However, their nomenclature was adopted from statistical analysis of microarray data in R^29^. The rationale for putting these files to the investigation level is to have unique sample and measured variable IDs across all studies. This convention can be adapted for multipartner investigations where each partner stores their work within its study and it thus makes sense to store phenodata files at the study level. For easier adaptation for wet lab researchers, pISA-tree assay metadata files allow for input of file paths or URLs for experimental protocols or data instead of providing full text.

pISA-tree users and data stewards have to be aware of some of its limitations, inherent to its design. One is the limitation of the maximum path length to 247 characters in Windows operating systems, which can only be circumvented by changing the Windows registry key. To avoid this issue, the usage of short but descriptive directory and file names is recommended. In addition, no automatic file and directory version control is enabled, except by using subsidiary software. In the user manual (Supplementary File 1), we advise the users on how to manage phenodata and featuredata file versioning, however for all other files a different versioning logic can be applied. The lack of version control can also be solved by uploading to FAIRDOMHub promptly after each significant update to the project. Taking into account the accessibility premise from the FAIR data principles, integration with FAIRDOMHub is also an elegant solution for sharing data with collaborators, stakeholders or the public.

Although pISA-tree was designed for small to medium sized projects, it has proven to be flexible enough to be adapted for larger projects with multiple independent research partners, which we showed in two ERACoBiotech projects SUSPHIRE (http://susphire.info/) and INDIE (https://indie.cebitec.uni-bielefeld.de/), both including five partners from three different countries and researchers with diverse fields of expertise, and H2020 RIA project ADAPT (https://adapt.univie.ac.at/) with 17 partners from 8 countries.

To conclude, we believe that pISA-tree will facilitate adoption of data management practices by individual researchers that will in long term lead to data FAIRness, adoption of open science principles and ultimately, new scientific discoveries.

## Methods

### Batch scripts

pISA-tree scripts and template files are organised in the main directory and the Templates subdirectory. The `Templates` subdirectory is further subdivided into `x.lib, DRY` and `WET`, which contain templates for step-by-step creation of bat files, and dry-lab and wet-lab assay classes. The batch script named `makeProject.bat` is stored in the main directory, and is the one to be used at the start of the pISA-tree structure creation (see manual in Supplementary file 1).

The main Windows executable batch file, `pISA.cmd`, is stored in the `\Templates\x.lib` directory. Depending on the user’s progress in building the pISA-tree structure for their project, this script generates additional `makeInvestigation.bat`, `makeStudy.bat` and `makeAssay.bat` at the appropriate levels within the project and initialises template metadata files. The directory names are automatically given a prefix defining the level i.e. “`_p_`” for project, “`_I_`” for investigation, “`_S_`” for study and “`_A_`” for assay, which facilitates visual inspection and general usage of the structure. In addition, assay directory names are automatically given an “-`<AssayType>`” suffix, that are defined in the `Templates` directory. The level-specific metadata files are text files containing a collection of tab-delimited key-value pairs in the form “`<key:>\t<value>`” and can be extended to suit the user’s needs via any text editor.

Another set of scripts can be used to inspect project directories and metadata files. The `showTree.bat` draws the pISA-tree directory tree structure to the `TREE.TXT` file, while the `showMetadata.bat` outputs a file containing all metadata in either plain text or markdown format, depending on the user’s position within the tree. The `xcheckMetadata.bat` highlights the missing values in the metadata files required by the minimum reporting standards, and kindly reminds the user to fill them in.

### Setting up the pISA-tree

To set up the pISA-tree, download the GitHub repository (https://github.com/NIB-SI/pISA-tree) into a local or network directory with read/write/execute permissions. This “pISA-tree root directory” ideally contains all pISA-tree projects. The setup does not require any administrative installation privileges of the user and the entire directory tree can be moved freely, if only relative paths are used in the metadata files, which is one of the recommendations given in the manual (Supplementary File 1).

### Modifying and extending assay types using metadata templates

The metadata associated with specific assay types is defined in template files in the `\Templates\DRY` and `\Templates\WET` class directories. Users can add new classes of assays, extend the existing types, or develop their own. To predefine the questions (and potentially multiple choice answers) that the user will be asked when creating a new assay, one adds rows of “<key:`>\t<value>`” pairs into the `<AssayType>_Template.txt` file. Here, keys define the questions and ‘/’ separated values provide menu choices. Users can also enter their own answers not provided in the template by selecting the option “`other`”.

Additional pISA-tree assay metadata templates are available at a separate, conceived as a community driven GitHub repository (https://github.com/NIB-SI/pISA-tree-assay-types) to be utilised for open science contribution of templates.

### Integration of pISA-tree with R and FAIRDOMHub

R package *pisar* enables repeatability of R-based analyses within the pISA-tree system. The users can retrieve recorded metadata information and the complete directory structure of the pISA-tree to reuse the dry-lab analysis protocols. The *pisa()* function retrieves the relative paths of the pISA-tree directory structure up to the project level whereas the *getMeta()* function retrieves specific fields (key values) from the metadata files. Thus, one can rerun R scripts without changing the code even if the metadata files are changed. As good practice, we recommend listing all metadata used at the end of the report.

The *seekr R* package was developed for communication of the pISA-tree with the SEEK API used by the FAIRDOMHub public repository and enables bulk upload of pISA-tree investigations. Currently, before uploading, the project page has to be created on the FAIRDOMHub web page to define user roles and permissions. The user can specify to exclude certain directories or files within the pISA-tree project directory tree by editing the “`seekignore.txt`” file. Further, uploading individual levels can be controlled by changing “`Upload to FAIRDOMHub`” key’s value in the level metadata file, which is, by default, set to “`Yes`”. In addition, the “`Sharing permission`” key sets the sharing of an individual level at FAIRDOMHub as either “`Private`” or “`Public`”, while the key “`License`” determines the license under which the data is shared to the public.

Bulk investigation upload to FAIRDOMHub using *seekr* is implemented with the R functions *skFilesToUpload()* and *skUploadFiles()*. The first function creates a list of files to be uploaded and the second function uses this list to prepare a JSON object for the specified investigation and all its accompanying files. It then and communicates with the SEEK API to create the corresponding FAIRDOMHub objects and then uploads the selected files. Package *seekr* uses the package *pisar* to obtain the local pISA-tree directory structure and metadata. A Jupyter Notebook is provided at seekr GitHub repository (https://github.com/NIB-SI/seekr) demonstrating the upload to FAIRDOMHub.

Unlike pISA-tree, FAIRDOMHub only allows linking files to the assay level. Therefore, to avoid having files disconnected from their corresponding levels, *skUploadFiles()* function creates a FAIRDOMHub study named “Investigation files” and a FAIRDOMHub assay named after the pISA-tree investigation where it stores the pISA-tree investigation metadata files. Likewise, it stores pISA-tree study metadata files in a FAIRDOMHub assay named after the study.

### pISA-tree metadata conversion to ISA-Tab

To enable interoperability with the ISA framework, we developed a converter from pISA-tree to ISA-Tab metadata format. The converter is implemented in the *pisar* package as *createISAtab()* function. It uses XML templates that contain information about the pISA-tree assay types that structurally correspond to the ISA-tools configuration files (v2015-07-02), and a mapping file that specifies how to convert fields from pISA-tree metadata files to ISA-Tab format. For each investigation, the function outputs ISA-Tab formatted metadata files that can be validated using ISAvalidator v1.6.5 from the ISA tools suite^13^. A Jupyter Notebook demonstrating straightforward conversion is available at pISA-tree-assay-types GitHub repository (https://github.com/NIB-SI/pISA-tree-assay-types).

## Glossary

ISA model: a metadata framework to manage an increasingly diverse set of life science, environmental and biomedical experiments that employ one or a combination of technologies. It is built around the investigation (the project context), study (a unit of research) and assay (analytical measurements) concepts^12^.
ISA level: common term for each of the concepts within the ISA model, either investigation, study or assay^30^.
ISA-Tab format: a hierarchical file format that captures the experimental metadata (https://isa-specs.readthedocs.io/en/latest/isatab.html) and is compatible with different spreadsheet-based formats for data sharing^13^.
ISA-JSON format: same as ISA-Tab but in JSON format instead of tabular.
Data standard: technical specifications or recorded agreements that include models, formats, reporting guidelines, and identifier schemas (https://fairsharing.org/).
Minimal information about experiment requirement: standards that describe minimal metadata that is required for a particular type of experiment to be FAIR (https://fairsharing.org/).
Data model: a model that organizes data elements and standardizes how the data elements relate to one another. A data model explicitly determines the structure of data (https://cedar.princeton.edu/understanding-data/what-data-model).
FAIR principles: guidelines to improve the Findability, Accessibility, Interoperability, and Reuse of digital assets^4^.

## Data availability

Example project data are available on FAIRDOMHub data repository under the project SEEK IDs 161 (https://fairdomhub.org/projects/161) and 252 (https://fairdomhub.org/projects/252).

## Code availability

The code and documentation for pISA-tree is available at http://github.com/NIB-SI/pISA-tree. R packages and documentation are available from https://github.com/NIB-SI/pisar and https://github.com/NIB-SI/seekr. The code is openly available under MIT license terms.

## Acknowledgements

The authors would like to thank Tjaša Lukan for testing pISA-tree, Valentina Levak and Katja Stare for helping with assay type template design, Wolfgang Müller for helpful advice and discussions, and Carissa Bleker for critical reading and language revision. This work was funded by the Slovenian Research Agency grants (P4-0165, J4-4165, J4-7636, J4-8228, J4-9302 and Z7-1888), BioPharm.Si project (OP20.00363), funded by the Ministry of Education, Science and Sport of Republic of Slovenia and European regional fund, European Union’s Horizon 2020 research and innovation programme projects ADAPT (grant agreement No. 862858) and EOSC-Life (grant agreement No. 824087), and European Research Area Cofund Action ‘ERACoBioTech’ (grant agreement No. 722361) projects SUSPHIRE and INDIE, which received funding from the Slovenian Ministry of Education, Science and Sport. For Fig. 2, logos of R, SEEK, FAIRDOMHub, and isatab were retrieved from https://www.r-project.org/logo/, https://fair-dom.org/about-fairdom, and https://isa-tools.org/software-suite.html, respectively.

## Author contributions

M.P. and M.Z. wrote the first draft and coordinated manuscript editing.

A.B. wrote the batch scripts and both R packages.

A.B., A.C., K.G., Š.B. and Ž.R. conceptualised the idea.

A.C., M.P., M.Z., K.G, Š.B. and Ž.R. wrote documentation and exhaustively tested the pISA-tree system.

M.P., M.Z., K.G, Š.B, A.C and Ž.R. contributed to the usability and improvement of the system.

K.G. secured funding, supervised and managed the project.

All authors read, reviewed and edited the manuscript draft and approved the final submission. All authors also contributed to the writing of a detailed user manual.

