## Supplementary File 1 for "pISA-tree - a data management framework for life science research projects using a standardised directory tree"

### FAIR DATA MANAGEMENT USING pISA-tree: STANDARD PROJECT DIRECTORY TREE

2022-05-26

https://github.com/NIB-SI/pISA-tree

### What is pISA-tree?

pISA-tree is a system for organisation of your experiments in the **F**indable, **A**ccessible, **I**nteroperable and **R**eusable (FAIR) manner - thus allowing integrative multiscale and multilevel analyses. It is set in accordance with ISA-tab standard and is compatible with FAIRDOM, using SEEK and JERM (Just Enough Results Model) frameworks as a basis.

pISA-tree provides a set of batch files (script files with extension .bat) that are used to create a standard directory tree for research projects. Batch scripts are executable on Microsoft Windows operating systems (OS) via Command Prompt (cmd). For Linux/Unix-like OS first install Wine. Command-line access in Wine is similar to Windows cmd and is invoked by typing wine cmd in the terminal.

Detailed instructions for installation are given in README.MD file on GitHub (https://github.com/NIB-SI/pISA).

### Definitions

**Sample** – material collected in the experiment

The definition of a sample is a complex matter. It starts with the sample collection itself (Fig.1).

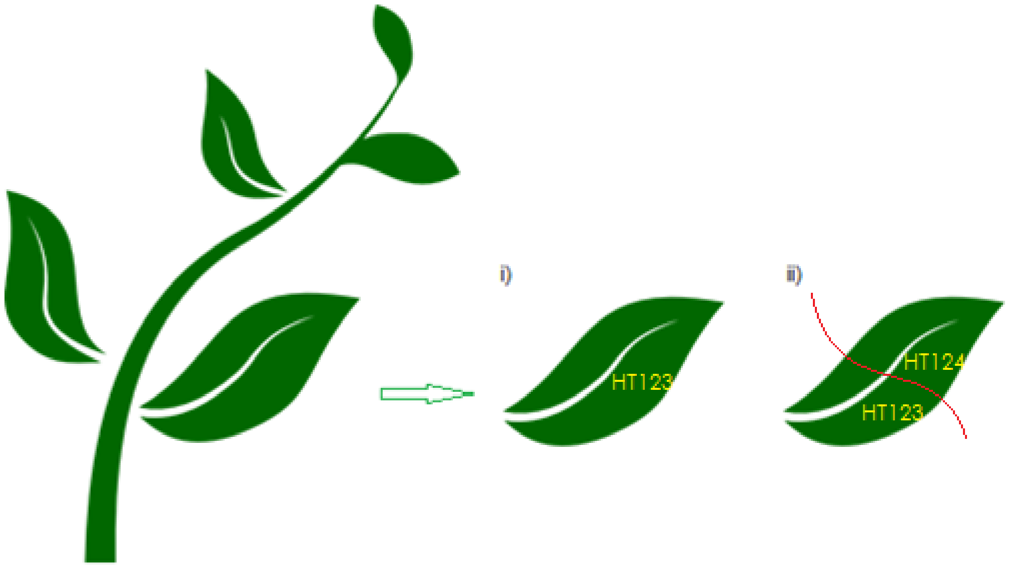

**Fig.1:** Sample collection example, showing assignment of sample IDs

i) If a full leaf would be homogenized and used in two techniques, transcriptomics and metabolomics, it would get the same ID (e.g. HT123).
ii) However, if the leaf would be first cut in half and then one part of the leaf would be used for transcriptomics and other part for metabolomics, leaf parts would get different identifiers (e.g. HT123 and HT124), meaning that one leaf may have more than one unique identifier, each of which identifies it for a different purpose. The system still has to allow us to extract the information that these two samples are from the same experiment, same plant and same leaf. This information is captured in phenodata file (see below).

**Analyte** – molecules extracted from the sample and analysed in an assay

Samples are further processed, e.g. RNA, DNA or proteins are isolated. If the same sample is used for isolation of different molecules or if something was wrong with procedure and the sample(s) need(s) to be processed again, then these **sample ID's** would be repeated several times and the traceability of our analysis would be lost (see Fig.2).

This is why we introduce the term **analyte**, for which we can create additional unique IDs that combine both the sample ID and the substance that was produced during analysis (e.g. HT123__RNA, HT123__cDNA, HT123__cDNA50x, see Fig.2). These additional analyte IDs are automatically created for wet lab assays of specific type (see subchapters Phenodata files and Assay).

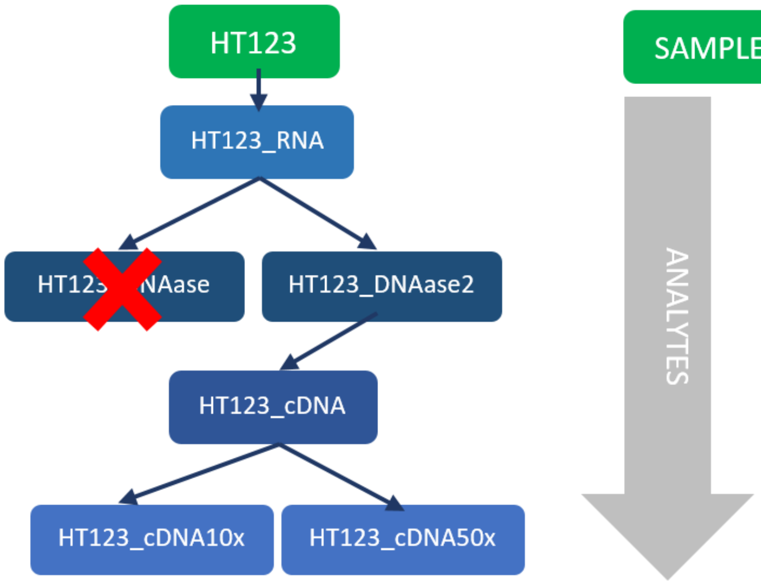

**Fig.2:** Example of Sample and Analyte IDs. From sample HT123, RNA was extracted, producing analyte HT123_RNA. This was treated with DNAse to produce HT123_DNAse. As the reaction was not successful, the step was repeated (HT123_DNAse2). The latter was used for dilutions (HT123_cDNA10x and HT123_cDNA50x), while HT123_DNAse was discarded.

**Phenodata file** – master sample file, document with sample descriptions (see Fig.3)

**Feature** – measured variable, e.g. pH, gene expression, protein abundance…

**Featuredata file** – list of features measured in the experiment, with some descriptions etc (see Fig.3)

**Sample ID** – an unique identifier determining a sample, usually in a short alphanumerical form e.g. HT123 (see Fig.2 and Fig.3)

**Analyte ID** – an unique identifier determining a substance analysed in the assay, usually in a short alphanumerical form e.g. HT123_RNA (see Fig.2)

**Feature ID** – a measured variable identifier, e.g. probe ID, gene ID, qPCR amplicon ID, … (see Fig.3)

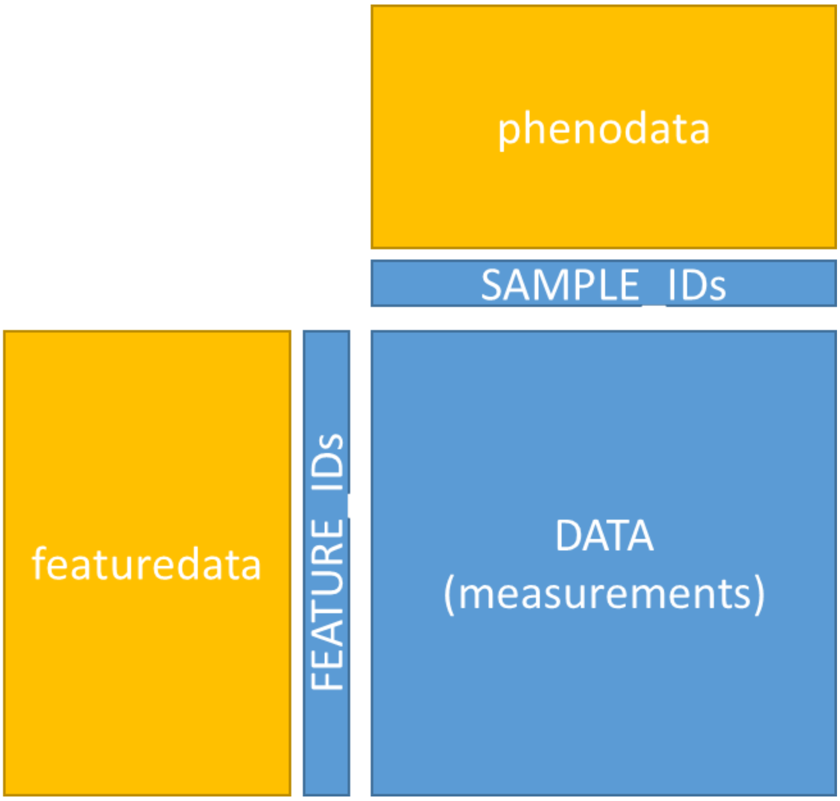

**Fig.3:** Schematic overview of relationship between phenodata and featuredata in high-throughput experiments.

**Minimimal information about experiment** – description of general information on how experiment (assay) was performed that provides us enough information for reproduction of experiment

**Metadata** – information describing the experiment and structured within different pISA-tree levels. Metadata about samples is encompassed in phenodata and metadata about features in featuredata. Metadata of assay should be structured according to recommended 'minimal information about experiment' standard.

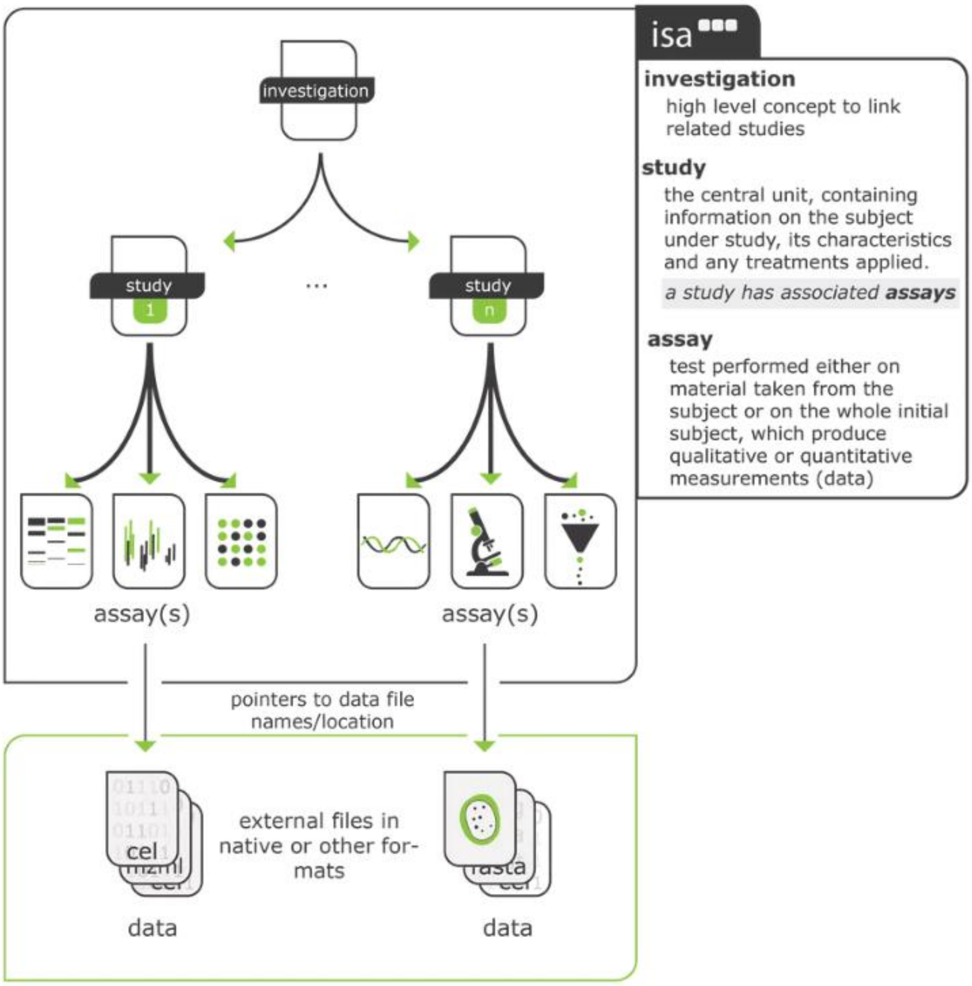

**Fig.4:** Schematic overview of the ISA-tab standard levels (Sansone *et al.,* Nat Genetics 2012) that are also used as levels of the pISA directory tree.

**ISA levels –** investigation/study/assay levels. A hierarchical data structure for linking and storing experimental data and metadata (see Fig.4).

**pISA-tree level ID –** a short name of project/investigation/study/assay levels; usually acronyms or abbreviations are used.

### Step-by-step instructions

To properly manage and annotate the data within the project one needs to design pISA project before the first samples are collected, that is when the experimental design is prepared.

#### pISA levels

You have to create a local folder (root directory), which will serve as the top pISA-tree level and will contain your future projects (on Fig.5 it is called **pISA_projects**). The root directory contains the **makeProject.bat** batch file whereas batch files for creating other levels are stored in the Templates folder (see Fig.5). Appropriate batch files are automatically copied into newly created levels.

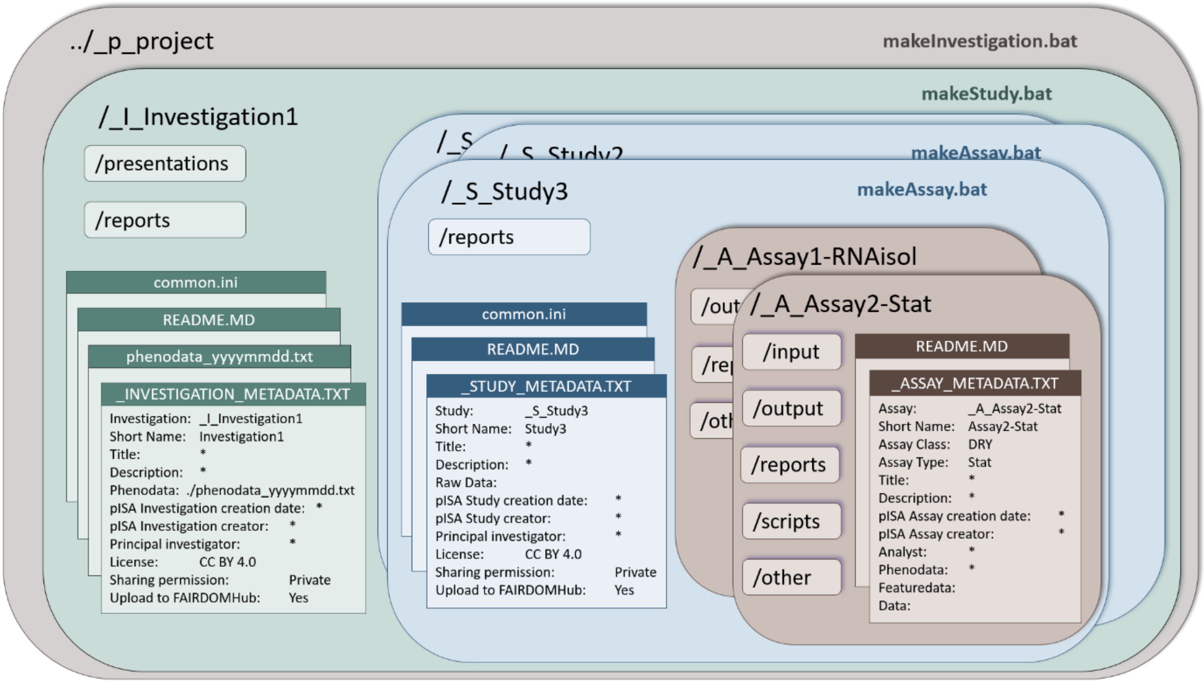

**Fig.5:** pISA-tree levels with corresponding subfolders and files that are automatically generated by batch scripts.

**Project** is organised as a collection of one or more **investigations**. An **investigation** is similarly organised as a collection of one or more **studies**. Each **study** has its own collection of one or more **assays**. Assays, either wet-lab or dry-lab, can be of specific type (e.g. MicroArray, NGS, Modelling, Statistical Analysis, ...) and are structured accordingly.

Here are some examples for each level of the pISA directory tree:

| pISA-tree | Description | Level ID examples |
|---|---|---|
| **Project** | | Lesions |
| **Investigation** | *The high-level concept to bring together related studies.*<br>Contains the Master sample table (named phenodata) | HT<br>*(stands for Hormonal treatments)* |
| **Study** | *The central level, containing information on the subject under study, its characteristics and any treatments applied.*<br>One biological experiment; e.g. batch of plants in the growth chamber, batch of fermentations performed in parallel, …<br>Exceptionally, a new study should be defined also when you are integrating data between studies. If the integration is within a study itself, then new assays are generated. | ser1_treat1_stu<br>ser1_ treat2_stu<br>ser2_treat1_ath |
| **Assay** | *Test performed either on material taken from the subject or on the overall initial subject, which produces qualitative or quantitative measurements (data).*<br>One test; it can be a batch of chips, qPCR plates…. This can be organised according to the researcher's preference. Wet-lab and dry-lab assays have different features to assist the researcher and consequently different structure. | PLT001<br>May18<br>ANOVA3<br>linRegM |

**Table 1:** project, Investigation, Study, and Assay levels definitions and examples.

#### Main batch files

- **makeproject.bat** - makes a new **project** directory tree
- **makeInvestigation.bat** - makes a new **investigation** directory tree (subdirectory tree within the **project**)
- **makeStudy.bat** - makes a new **study** (subdirectory tree within the **investigation**)
- **makeAssay.bat** - makes a new **assay** (subdirectory tree within the **study**)

#### Creation of the directory tree

The directory tree is a way to enforce the subordination of pISA levels. To emphasize the level type, directory names are constructed automatically using the standard prefix and short level ID. Standard prefixes are:

- _p_ for project
- _I_ for investigation
- _S_ for study
- _A_ for assay

##### project

To create a new project, run (double click) the file **makeProject.bat** and enter the project ID (short project name without spaces and special characters; see Annex 2). This will make a directory tree, metadata files and a local copy of **makeInvestigation.bat**. Short project name (ID), automatically prefixed with _p_, is used as the name of the directory. For example, if you set the project ID as blah the project directory name will automatically become _p_blah.

*Note: If a Windows Defender warning message appears while executing batch files, click 'More info' and then 'Run anyway'. You have to do this only once for each batch file.*

#### Investigation

To create a new investigation, run the file **makeInvestigation.bat** and enter the investigation ID (short investigation name without spaces and special characters; see Annex 2). This will make a directory tree, metadata files and a local copy of **makeStudy.bat**. Short investigation name (ID), automatically prefixed with _I_, is used as the name of the directory. The investigation directory name for the investigation bleh will be _I_bleh.

#### Study

To create a new study, run the **makeStudy.bat** and enter the study ID (short study name without spaces and special characters; see Annex 2). This will make a directory tree with several standard folders, metadata and auxiliary files and a local copy of **makeAssay.bat**. The study folder name will be _S_bloh for a study with short name (ID) bloh.

*Note: New study should be initiated for each new batch of samples that is going to be collected!*

#### Assay

Analyses for each study are stored in the folder of that study. To make a new assay, run the **makeAssay.bat** file.

First, you will be asked to choose between the assay wet- or dry-lab **Class**:

- Wet-lab e.g. measurements on the biological material (MicroArray, RNA-seq, qPCR, ...)
- Dry-lab e.g. process data (Statistics, Modelling, data integration, ... )

Second, you will enter the assay **Type** (i.e. RNAisol, qPCR, RNA-seq, GC-MS) and assay ID (Short name, for example RNA1). Short assay name and type (separated by '-' and prefixed by _A_) are used as the name of the assay directory tree (for example: _A_RNA1-RNAisol). The structure of subfolders and automatically generated files in shown in Fig.5.

*Note: defining the assay type can bring in some additional item descriptors that can help you with data management of particular experiment. Each assay type, however, has to be carefully designed. For wet-lab, RNA isolation coupled with DNase treatment and reverse transcription (RNAisol) is available. The user can manually create new assay types as explained in Annex 3 or create them on the fly by selecting Other from the batch script menu.*

Third, you will be is asked to choose the phenodata file that will be used in the particular assay.

*Note: phenodata file must be in text format, otherwise **Analytes.txt** file will not be generated. It is recommended that you enter the Assay ID in Phenodata first, so Analytes file will be generated automatically (see Phenodata).*

When creating either of these levels a certain folder structure is created. Descriptions of generated subfolders and required files are given in the table below:

| | pISA-tree levels | | | |
|---|---|---|---|---|
| **created directories** ***/ files*** | **INVESTIGATION** | **STUDY** | **ASSAY** | |
| | | | **wet-lab ASSAY** | **dry-lab ASSAY** |
| *level-specific metadata files* | _INVESTIGATION_METADATA.TXT<br><br>phenodata_yyyymmdd.txt *(master sample table)*<br><br>featuredata_yyyymmdd.txt *(variable description file)* | _STUDY_METADATA.txt | _ASSAY_METADATA.TXT<br><br>analytes.txt *(analyte table)* | _ASSAY_METADATA.TXT<br><br>analytes.txt *(analyte table)* |
| /input | - | - | - | *input data set(s) or file(s) containing path(s) to input data* |
| /scripts | - | - | - | *scripts, pipelines, workflows or commands used for the analyses* |
| /output | - | - | *pre-processed results* | *final results* |
| /output/raw | - | - | *raw result files, e.g. data collected from the machine* | - |
| /reports | *summary reports for project partners and stakeholders* | *sample handling or any other study related protocols or reports* | *assay specific protocols, laboratory logs* | *documented procedures, protocols, log files with tool(s) version(s) used for the analyses* |
| /presentations | *any presentation related to the project* | - | - | - |
| /other | - | - | *other relevant files not included elsewhere* | *other relevant files not included elsewhere* |

**Table 2:** Automatically generated pISA-tree directories and metadata files and recommended allocation of data files. Each level contains level-specific metadata files and directories containing data. '-' directory not created at this level.

### Metadata files

Several metadata files need to be prepared by users or are automatically generated by pISA-tree.

#### pISA level metadata files

Each level has a *_LEVEL_METADATA.TXT* file, a file with additional information needed to describe the experiment with enough information to be reproducible. This metadata files are tabulator-delimited text that list informative items for specific pISA levels in two columns:

1. item name (ended by a colon)
2. item value

Item value can be some text, for example investigator's name or longer study description, analysis description etc., or path to the phenodata file. Each item pair in the metadata should be typed in one line. Be careful if the metadata contains prime symbol (', as in 5'), it is better to spell it out, like 5-prime. For other unfavoured characters see Annex 2.

Two examples of the metadata entries are given below (tab character is shown as right arrow →):

Investigator: →Miha Mihav
Phenodata: →./phenodata_20181010.txt

When starting new project, investigation, study or assay, pISA-tree will guide you through the questionnaire to collect the required metadata.

#### README.MD files

At each pISA level a dummy **readme.md** file is automatically generated. These are free-form text files and can be used to make notes that explain the content of the directory, changes made to files etc.

#### Common.ini files

When running batch file to create a new project, investigation, study or assay, the user is asked to enter basic metadata (as described above). Some metadata are however identical for all studies and assays within one investigation (or similar for other pISA-tree levels). To avoid multiple entry of these metadata with every new study, user can enter such information into **common.ini** file. This file is created as a dummy file in pISA-tree root directory and will be automatically copied to newly created pISA-tree levels. This file contains following content:

Principal investigator:→*
License:→Creative Commons Attribution 4.0
Sharing permission:→Private
Upload to FAIRDOMHub:→Yes

The last three lines are related to synchronisation with FAIRDOMHub and need to be filled in if you plan to synchronise your pISA-tree with FAIRDOMHub automatically. License options are listed here: https://docs.seek4science.org/help/user-guide/licenses.html.

The **common.ini** files should be modified by the user to enter fields and metadata that are fixed for a particular project/investigation/study/assay (e.g. the principal investigator name, contact address, etc). Information in these files will be automatically appended to metadata files for all subordinated pISA-tree levels.

*Note: In computer sciences *.ini files usually contain initial values and settings, thus here this file extension is used.*

**_LEVEL_METADATA.txt** files and **common.ini** file are plain text files. You can open and edit them with any text editor (e.g. Notepad++, WordPad, ConTEXT, Nano, …), Excel, OpenOffice Calc, … at any time (not just when starting a new level in pISA-tree). In some text editors the tab character that is separating item names and item values might be invisible. You can visualise it by enabling the "show symbols" or "show all characters" option. If you use Excel, the file will be presented in two columns and might be more readable and easier to edit. In this case, do not forget to save the file opened in Excel as Text (Tab delimited) file and do not change its name nor extension (.txt or .ini).

#### Phenodata files

Phenodata files (the name of the file originates from the golden age of microarrays) are tabulator delimited text files that describe your **samples**. Sometimes they are also referred to as Master sample files. Phenodata files are created so that they contain date of creation (e.g. phenodata_20181010. txt; see Note2) and are **stored in the Investigation folder**. Every start of new Study is related to the collection of new samples in wet lab. Already before starting the real experiment (e.g. growing plants), one should create a phenodata file together with the basic pISA-tree structure.

If you use Excel, the file will be presented in columns and might be more readable and easier to edit. In this case, do not forget to save the file opened in Excel as Text (Tab delimited) file and do not change its name or extension (.txt). Excel file should have only one sheet. Remove all filters before saving as TXT. Column headers of the phenodata file are partially prescribed, but any additional columns that might help to better describe collected samples can be added.

All samples used in an investigation must have unique sample IDs (unique keys), which are a combination of the two-letter study acronym (e.g. HT for hormonal treatments) and a three-digit number, e.g. HT001–HT999. By definition, unique sample ID means that within the same phenodata file there will not be two distinct rows that have the same values of sample ID.

Besides sample ID, which is always in the first column, phenodata file should contain Sample Name (longer and more descriptive). Further columns should contain sample descriptions, like for example: time after start in days (1, 2, 3, ...), treatment data (mock, PVY, …), genotype (NT, coi1, NahG, ...), position of the sample on the plant (upper leaf, ...) and any further information you consider relevant for analyses and reproducibility. When creating these descriptions, you should not use any spaces nor special characters (see Annex 2). Times and Dates should have consistent formatting within the same Phenodata file. Column names should always be unique. Under column name corresponding to the Assay ID (e.g. RNA1-RNAisol) please mark which samples will be analysed in this assay (e.g. YES, X, 1, ...) and leave empty for those not analysed. Names of the Assay should be written with full name (see Note 3). **Analytes.txt** file will be generated automatically for selected samples within Assay directory.

An empty phenodata file is automatically generated when you create a new pISA-tree investigation.

*Note1: to allow computer readability of phenodata, allowing for easy automatic integration of results, standard vocabulary should be used when filling in the phenodata file. Standard vocabulary means that you always use the same word for the same description. The 'same word' here means to be consistent using the uppercase/lowercase combinations, hyphens, underscores and other appropriate characters. Information on minimal information to be entered can be found in various standards (see Annex 1).*

*Note2: sometimes* ***corrections of the phenodata files*** *are needed; for example, when a sample is misannotated or is misclassified. To allow traceability of all dry lab analyses, when the phenodata is changed, it should be saved with the new date. In this way we can trace which phenodata file was used for which assay and repeat the analysis, if necessary. Note that opening a new study and adding new samples to the phenodata is not considered as a correction.*

*Note3: for any type of analyses, especially dry-lab assays, it is very practical to add info in which assays which samples should be used. Prior to the creation of the assay level (using* **makeAssay.bat***) the user should add a column (to the right of the last existing column) into the phenodata file marking the samples that will be analysed in that particular assay. The column name should be Assay name of the planned assay and the samples should be marked by adding a single character into the corresponding cell.*

*An additional feature is offered in pISA-tree helping you in analyses/reporting on the level of assays. When* **makeAssay.bat** *file is executed, user will be asked to enter the standard and assay specific items (metadata). The* **Analytes.txt** *file (see example in the table below) will be generated at the Assay level combining the information about the samples in the phenodata file and information entered as metadata:*

| SampleID | Sample Name | Homogenisation_ protocol | Operator | Date_Homo genisation | RNA_ID | ng/ul | ... |
|---|---|---|---|---|---|---|---|
| SMP_001 | Leaf 1 | Rneasy_Plant | Bob | 24. 04. 2018 | RNA_SMP_001 | | ... |
| SMP_003 | Leaf 3 | Rneasy_Plant | Bob | 24. 04. 2018 | RNA_SMP_003 | | .. |
| SMP_007 | Leaf 7 | Rneasy_Plant | Bob | 24. 04. 2018 | RNA_SMP_007 | | .. |

**Table 3:** Example of Analytes.txt file for the RNA isolation Assay. Columns 1-6 were generated automatically by the script, whereas the other content (results of the analysis) has to be entered by the user.

#### Featuredata files

Featuredata file (i.e. annotation file) lists and describes the features (e.g. gene, metabolite) measured in a particular assay (biological experiment). Besides the unique IDs (e.g. geneID, metaboliteID, ...), the file that describes the features also provides additional information about that features (e.g. short name, description, Gene Ontology terms, EC, MapMan Bin, ...), any technical issue (e.g. specificity problem, quantification problems), etc.

The file should be created or downloaded (**.gal** file in the case of microarrays, **.gff** file in case of RNASeq, ...). The file should be prepared in a tab delimited format where the first column contains list of all features and is named *featureID* (see also Note 3 below), followed by any number of columns that give improved knowledge and understanding of the feature.

*Note 1: Although the annotation files can be quite complex, they have to contain at least two columns: featureID and Description.*

*Note 2: For microarray analysis this file normally includes also information on feature positions on the microarray which are provided by the manufacturer of the microarrays.*

*Note 3: For all transcriptomics (microarrays, NGS, qPCR) and proteomics experiments we will link the features to corresponding genes. Consequently, first column in the Annotation file should list GeneIDs and be named "geneID".*

#### Auxiliary batch files

These files can help you with your data management issues but are not obligatory for FAIRness of your data:

##### showMetadata.bat

Collects all metadata files in a tree below the current level. Descriptions are typeset in either **METADATA.TXT** (plain text file) or **METADATA.MD** (plain text file in a markdown format; all text files can be edited by any text editor, e.g. Notepad, Wordpad or Excel and Word as long as they are saved as the text files. Use 'Open with' option to select the non-default program to open such data).

##### xcheckMetadata.bat

Checks all metadata files for missing required information (*) in a tree below the current level. Produces the file named **xCheckMetadata.md** which is similar to the one produced by showMetadata.bat but lists only lines with asterisks (*).

##### showTree.bat

List a directory tree below the current level in the file **TREE.TXT**.

##### update.bat

Replaces batch files in existing folder tree (all existing projects, investigations, studies and assays) with the updated versions from the root and Templates/x.lib subdirectory. After downloading an updated version of pISA-tree from GitHub, extract and replace all files in root and Templates directory. Run the update.bat file to update all batch files in all existing levels.

### Annex 1: Standards helping in setting up appropriate metadata files

Plenty of various platform dependent standards exist for the description of experimental data; consequently, all these standards are assay dependent (e.g. qPCR assay that involves sample preparation for it).

- ISA-TAB creator allows us to modify existing templates to suit our purposes or create new ones
- some of the existing templates: default ISA-TAB templates, MIAPPE template that improves on the phenotyping, for metabolomics: CIMR-MTBLS, MetaboLights; ScientificData templates
- pISA tree templates are stored in folder /Templates/WET/

- relevant minimum information standards (more info on FAIRsharing webpage):

| Standard | ExperimentType | Description |
|---|---|---|
| **MINSEQE** | sequence reads | minimum information about a high throughput sequencing experiment |
| **CIMR** | metabolomics | core information for metabolomics; see MIAPE-MS too |
| **MIQE** | transcriptomics | minimum information for qPCR experiments |
| **MIASE** | models | minimum information about a simulation experiment |
| **MIRIAM** | models | minimum information required in the annotation of models |
| **MIAPA** | sequences | minimum information about a phylogenetic analysis |
| **MIACA** | - | minimum information about a cellular assay; high-throughput cell biological analyses (cells in culture); extension of minimum information captured by primary nucleotide sequence archives |
| **MINI** | electrophysiology | minimum information about a neuroscience investigation; electrophysiology |
| **STRENDA** | - | standards for reporting enzymology data guidelines |
| **MIAPPE** | Phenotype features | Minimum information about plant phenotyping |

### Annex 2: Characters to avoid in directories, filenames, tables and IDs

#### Do not use any of these common illegal characters/symbols:

- # pound (hashtag)
- < left angle bracket
- $ dollar sign
- + plus sign
- % percent
- > right angle bracket
- ! exclamation point
- ^ circumflex accent
- & ampersand
- * asterisk
- ‘ single quotes
- | pipe
- { left bracket
- ? question mark
- “ double quotes
- = equal sign
- } right bracket
- / forward slash
- : colon
- \ backslash
- blank spaces
- @ at sign

#### Avoid naming files using prefixes designated for project/Investigation/Study/Assay levels:

- _p_
- _I_
- _S_
- _A_

#### Keep these rules in mind:

1. do not start or end your filename with a space, period, hyphen, or underline.
2. keep your filenames to a reasonable length
3. operating systems are case sensitive
4. avoid spaces and special characters (e.g. + and − are mathematical symbols).
5. do not use data types and keywords for table or column names, also do not pick names that will change meaning

#### Appropriate folder names:

- myDocuments
- my_Documents
- MyDocuments

### Annex 3: Developer tools

Here some additional features of pISA-tree app are listed which are not applicable for the standard user, but more for the ones that would like to extend it.

#### Adding new Assay Classes and Types

Subdirectories within assay directory trees, for different Classes, differ slightly, according to the need of the specific **Class**. Assay classes and types are defined as subdirectories of the Templates directory. An example is "../Templates/Wet/RNAisol". For this example, the directory name defines the assay **Class** as "Wet" and the subdirectory name assay **Type** as "RNAisol". To add another class, create directory *myclass* within Template directory: "../Templates/myclass". To add another type of, for example *Wet*-lab assay (here named *mytype*), create it on the fly by selecting Other from the batch script menu or create a new subdirectory within appropriate Class directory with the name *mytype*: "../Templates/Wet/mytype".

In addition to the basic items, one can also use assay specific items, depending on the assay type. The assay specific items are pre-specified in the **meta_AType_Template.txt** and **Analytes_Template.txt** files, placed within the appropriate *Class/Type* subdirectories. The **makeAssay.bat** batch file will accordingly generate questions (if any) to add information to assay metadata file. The **meta_AType_Template.txt** and **Analytes_Template.txt** files are specific for each used assay Type in your system.

The **meta_AType_Template.txt** and **Analytes_Template.txt** files are plain text files. Each line represents one "Item name - Item value" pair, separated by the tabulator character (illustrated below as the right arrow →). The first of the pair – "Item name", will appear as the assay specific question during the assay creation. The second of the pair – "Item value", will be either offered or has to be entered manually.

An example of the **Analytes_Template.txt** file:

***Item name → Item value***
Isolation Protocol →Rneasy_Plant/ZymoRNA
Operator →John/Bob/Katja/Anna
Date Homogenisation→%today%
RNA ID → RNA_$
ng/ml → Blank

The **meta_AType_Template.txt** or **Analytes_Template.txt** will not need to be tackled with by standard user of pISA-tree application.

The user will be asked about the assay specific items (defined by assay specific **Analytes_Template.txt** file) when running **makeAssay.bat** and those will be included in the **_ASSAY_METADATA** file. In addition, they are used as the assay specific description of samples used in an assay and are automatically added as the assay specific extension to the phenodata file. Assay specific metadata will be copied into columns of the **Analytes.txt** file, which contains information about the samples used in the assay.

Syntax rules in item value part are used for support of choices in menu-like data entry. This reduces errors in spelling, spacing, and use of the character case.

**Fields with one or more choices**

Item value choices, if more than one, are separated by the slash (/) character. See the example above for the items named Isolation Protocol and Operator. To select the operator name, a simple menu will be presented to the user:

1 John
2 Bob
3 Katja
4 Anna
5 Other

User will use numbers (1 to 5) to select the name to use. The last line ("Other") is automatically added and enables ad-hoc addition of any new choice. If the choice is likely to occur in future, it can be added into the **analytes.ini** file.

##### Date field

Date fields are considered in the same way as ordinary choice fields. Special bookmark %today% will be replaced by current date in a data entry menu.

##### Sample ID replacement

New sample related identification codes are sometimes needed. Sample ID can be automatically inserted in the place of a dollar character ($) to form new IDs. In example above for the field RNA ID and Sample names HT123, HT124, HT125 one would get new IDs: HT123_RNA, HT124_RNA and HT125_RNA.

##### Blank fields

The word Blank as item value signals the column that has to be left blank in the Analytes.txt file.

| | File name | Add/edit items | Entry | Used for | Included in: |
|---|---|---|---|---|---|
| Assay type specific | **meta_AType_Template.txt** in AssayType folder | Yes | Typed during creation | Assays of this type | Metadata file |
| Analytes specific | **Analytes_Template.txt** in AssayType folder | Yes | Typed during creation | Assays of this type | Metadata file and Analytes.txt |

#### Metadata-only specific fields

Fields within **meta_AType_Template.txt** files starting with '+' sign will not be shown when running **makeAssay.bat** script, however they will be added to the corresponding metadata file. In provided templates, these fields are used to define Measurement Type and Technology Type name/value pairs, and it is considered good practice to write them at the beginning of the template file.

### Additional notes

##### Additional Note 1

You can rename directory on all levels (project, Investigation, ...).
An example:

rename the project directory name, and

in *_PROJECT_METADATA.TXT* file, change project ID and short name to desired one.

### Notes on distributed assay type templates

#### RNAisol

This wet-lab template covers the wet-lab procedure of sample collection, storage, RNA isolation, DNase treatment and cDNA preparation. This is a typical molecular biology workflow preceding other analysis such as PCR, qPCR, RNA-Seq, microarrays or cDNA cloning

**Instructions on RNAisol assay specific data and metadata storage**

Store Nanodrop and/or Bioanalyzer file exports as well as agarose gel images in assay directory "output/raw/". Any files derived from the raw files e.g. Excel files combining several raw files or annotated gel images should be stored in assay directory "output".

**ASSAY METADATA**

We recommend populating this list when creating the assay, however certain metadata can be added or modified later (if so, the assay's ANALYTES.TXT file should be modified accordingly).

**Assay-specific metadata explanations**

RNA ID – the suffix for the RNA sample ID e.g. "_RNA"

Homogenisation protocol - choose between fastPrep, TissueLyser, mortar or Other

Date Homogenisation - date of tissue homogenisation

Isolation Protocol – choose between Rneasy_Plant, ZymoRNA or Other

Date Isolation - date of RNA isolation

Storage RNA – the place where the RNA is stored such as the freezer box ID e.g. CU0369

DNase treatment protocol – shortly describe the protocol or link to protocol file

DNase ID – the suffix for the DNase treated RNA sample ID e.g. "_ DNase"

Date DNAse_treatment - date of DNAse treatment

Storage_DNAse_treated - the place where the DNase treated RNA is stored such as the freezer box ID e.g. CU0369

Operator – name or acronym of the person doing the wet lab work

cDNA ID – the suffix for the cDNA sample ID e.g. "_ cDNA"

DateRT – date of reverse transcription reaction

**ANALYTES.TXT**

This file serves as a file for storing metadata (IDs of analytes from the assay) and the performed quantity and quality measurements. If it is generated, open the file in Excel and copy the following measurements into the following empty columns:

ng/ul – copy concentration measurements from e.g. Nanodrop or Bioanalyzer

260/280 – copy Nanodrop QC ratio measurements

260/230 - copy Nanodrop QC ratio measurements

### qPCR

Quantitative PCR (qPCR) is a standard technique for quantifying the amount of nucleic acids in the sample. It is a well-established method for determining specific gene expression changes using the relative quantification relying on stably expressed reference genes. It is often used to validate high-throughput transcriptomics methods e.g. microarrays or RNA-Seq.

**Instructions on qPCR assay data and metadata storage**

When creating a qPCR assay you should save the files that you export from the qPCR machine into the "output/raw" directory. Files exported from software for analysis of qPCR results e.g. quantGenius (quantgenius.nib.si/) should be saved into the "output" directory where you can also store your additional analyses e.g. graphs or combined exports from qPCR analysis software (use file name suffix "_compiled"). In the "reports" directory you should put presentation or log files describing the analysis workflow or file history.

**ASSAY METADATA**

The template was prepared by taking into account MIQE précis minimal standard guidelines for fluorescence-based quantitative real-time PCR experiments.

**Assay-specific metadata explanations**

Lab manager – name or acronym of the laboratory manager

#Assay optimisation/validation

Assay chemistry – choose the qPCR chemistry based on the design of amplicons. If you used different chemistries in the same assay, choose "Other" and select and paste all used chemistries

Assay optimisation protocol – file path, link or citation to the protocol for optimizing amplicon (can be one file for all parts of protocol)

Pippeting – choose between liquid handling station (pipetting robot) or manual pipetting

Reaction volume – final qPCR reaction volume in ul

#qPCR

qPCR platform – choose the platform (machine) used in the assay. If you used different platforms in the same assay, choose “Other” and select and paste all used platforms.

qPCR protocol – file path, link or citation to the protocol for qPCR (can be one file for all parts of protocol)

One-step RT-PCR (RNA as input) – choose if you performed reverse transcription and PCR in one step

Well format – choose well plate format

MasterMix – choose the qPCR MasterMix used in the assay. If you used different MasterMixes in the same assay, choose “Other” and select and paste all used MasterMixes

No. sample technical replicates – number of technical replicates of same reaction

No. sample dilutions – number of sample dilutions for each sample (used to determine PCR efficiency of each sample)

PCR controls used – choose controls used in the assay. If you used more than one, choose “Other” and select and paste them.

No. standard curve dilutions (if using) – number of dilutions of the sample (or sample pool) used for the standard curve in relative standard curve qPCR quantification method

Operator – name or acronym of the person doing the wet lab work

#Data analysis

Cq value determination software (if different from PCR platform) – software used to determine Cq values from the fluorescence signal if it was different from the one specified for the “PCR platform”.

Quantification software – software used quantification of nucleic acids in the samples from the Cq values

qPCR quantification method – choose the method or use “Other” to specify your own

Notes – any relevant notes

### LUM

The luminescence (LUM) assay is used to test gene promoter activity. When a promoter is active, the luciferase reporter gene is transcribed and after translation of mRNA to protein, one mature luciferase enzyme produces one photon of visible light. With the luminescence assay, promoter activity can be followed *in vivo* for a long period of time (up to several days).

**An example of luminescence assay setup for testing promoter activity**

A plasmid with luciferase reporter system is agroinfiltrated into tobacco leaves. Few days later, we excise leaf disks and put them in a 96 well plate - one leaf disk per well. Cut two disks in near proximity on the same leaf (“sampling in pairs”) and use one of them for control and another for treatment. This way, we don’t get information on promoter activity only as a ratio between average of all treatments and average of all controls, but also as a ratio between treatment and control of each leaf disk pair. This is important if we want to see how the ratio varies among different parts of the leaf, different leaves and different plants.

**<u>Instructions on how to store the data and metadata of above LUM assay setup in pISA-tree</u>**

<u>STUDY</u>: Name the STUDY based on the promoter you are studying e.g. CPI8, MC, PR1b. In case there are different versions of this promoter available (homologues), **all of them should be kept in the same STUDY**. For example, different versions of CPI8 promoter amplified from a cv. Rywal plant (CPI8.Ry1, CPI8.Ry14) and from a cv. Désirée plant (CPI8.De1, CPI8.De2, CPI8.De7) should be stored in the same Study directory named "_S_CPI8".

**<u>ASSAY METADATA</u>**

One luminescence experiment corresponds to one 96- (or 384-) sample well plate and one Assay directory. The name of the ASSAY should be the consecutive number of the LUM assay in the Investigation folder (check the **phenodata.txt** file before you make a new Assay). For example, for the first assay named "LUM1" the Assay ID would be "_A_LUM1-LUM". You should describe the assay more precisely using the Assay title (e.g. "LUM1-CPI8.De1-CPI8.De7") and description item values in the **_ASSAY_METADATA.txt** file.

If you test 2 different promoters (e.g. CPI8 and MC) in the same well plate, you should create an ASSAY directory in both STUDY directories (e.g. CPI8 and MC). Keep all data in the first ASSAY directory, while in the second ASSAY directory you should add the following statement to the "raw data" item in the **_ASSAY_METADATA.txt** file: "All data is kept in ../_A_LUM1-LUM". In the second ASSAY output directory, you might also create a shortcut to the results file in the first ASSAY directory.

**Assay-specific metadata explanations**

Lab manager – name or acronym of the laboratory manager

Operator – name or acronym of the person doing the wet lab work

Link to SynBio DM – link to SynBio data management file

Link to protocols – link to protocol file

Additional Notes – free text notes

Plate ID – well plate ID in the format PS### (# represent digits)

DNA introduction system – choose between agrobacteria mediated transient transformation, protoplast transfection, stable transformation or Other

SynBio GenotypeID(s) – the genotype ID from the SynBio data management file

Date of transformation/transfection/micropropagation -

Treatment – choose between the listed substances or Other and enter treatment

Date of treatment – date of plant treatment

Date of sampling – date of leaf disk sampling

Date of luminescence assay start – choose the current date (1) or Other and enter date

Time of luminescence assay start – choose the current time (1) or Other and enter date

Luminometer - choose between BioTek Synergy Mx or Other and enter machine name

Protocol (luminometer setup) - choose between Long Kinetic Experiment or Other and enter protocol name

Time [min] between two measurements (luminometer setup) – most common setup options are given to choose from, otherwise choose Other and enter it

Sensitivity (luminometer setup) - most common setup options are given to choose from, otherwise choose Other and enter it

Integration time [ms] (luminometer setup) - most common setup options are given to choose from, otherwise choose Other and enter it

Temperature (luminometer) at start of the experiment [degrees Celsius] - most common setup options are given to choose from, otherwise choose Other and enter it

Temperature (luminometer) at the end of the experiment [degrees Celsius] - most common setup options are given to choose from, otherwise choose Other and enter it

Time between start of sampling and start of experiment [min] - most common setup options are given to choose from, otherwise choose Other and enter it

Final concentration of luciferin [uM] - most common setup options are given to choose from, otherwise choose Other and enter

Silencing suppressor added - choose between none, p19 or Other and enter silencer suppressor name

**SAMPLE**

Each well with a leaf disk represents a sample and should get a unique ID (for all samples in Assays that are under the same Investigation). This includes blanks (non-agroinfiltrated leaf disks or empty wells used to measure background effects during measurement), controls and treated samples. This should be clearly noted in the **phenodata.txt** file in the appropriate column. The sample ID of each leaf disk should look like "PS001-01", where the first two letters indicate the Investigation acronym (in this case PS stands for Promoter Studies), the following 3 numbers indicate the Assay number and the 2-3 numbers after dash (-) indicate the individual well in the plate (1-96 in case of 96-well plates or 1-384 in case of 384-well plates).

**PHENODATA FILE**

First part of the sample ID (before dash) corresponds to PlateID in the **phenodata.txt** file and the second part (after dash) to WellID. Explanation of columns in the **phenodata.txt** file:

Promoter – promoter's name and version e.g. CPI8.De7

SampleWell – well position on the plate e.g. a1-h12

PartOfLeaf – choose between options: middle, margin, base, apex, base margin, apex margin, NA (NA stands for not available)

SampleName – we recommend the notation similar to the following: "p1l1_1", where "p1" stands for plant 1, "l1" for leaf 1 and "_1" for disk 1

CommentsOnPlantMaterial – comments such as "the leaf was damaged before sampling"

Treatment – choose between no, *in vivo*, *in vitro* (whether the plant was treated before sampling or leaf disks are treated *in vitro* simultaneously with the experiment)

InVivoTreatmentTime – time between the treatment and sampling if you performed *in vivo* treatment, otherwise leave empty

TreatmentWith – name treatment substance or condition e.g. JA, MeJA, SA, INA, Ethylene, ACC, Dexamethasone, heat, cold, wound and positive control, negative control or blank for controls

HormoneConcentration – concentration in mol/l or g/l. Mark the correct unit with "1" in one of the two following columns

TreatmentOtherQuantity – if you used any other physical quantity as a treatment, write its unit here

SampledInPairWithSampleWell – if you sampled in pairs, write the plate well position of the corresponding sample

Comments – observations made for leaf disks e. g. submerged or yellow after experiment, non-infiltrated by mistake (when realized after the assay), assay outlier etc.

Result – other notes

### GCMS

Gas chromatography mass spectrometry (GCMS) assay is used to identify and quantify different metabolites within a sample. This template covers the wet-lab procedure of sample collection, storage, preparation and analysis.

**ASSAY METADATA**

**Assay-specific metadata explanations**

Lab manager– name or acronym of the laboratory manager

Sample collection protocol - shortly describe the protocol or link to protocol file

Extraction protocol - shortly describe the protocol or link to protocol file

Chromatography protocol - shortly describe the protocol or link to protocol file

Mass spectrometry protocol - shortly describe the protocol or link to protocol file

**ANALYTES.TXT**

#Extraction

Extract ID – the suffix for the extract sample ID e.g. "_extr"

Extract Name – name of the extract sample

Extraction Method – short name of the extraction method

Date Extraction – date of performing the extraction

#Post Extraction

Derivatization or Labelling – choose between sylilation, acetylation, alkylation, none or select Other and enter it

Date Derivatization or Labelling – date of performing the derivatization or labelling

Derivatized or labeled Extract ID – the suffix for the derivatized or labeled extract sample ID e.g. "_extrD"

Derivatized or labeled Extract Name – name of derivatized or labeled extract sample

Other Post Extraction Procedures – any procedure performed after extraction except derivatization

Storage – storage of extract samples such as the freezer box ID e.g. CU0369

#Chromatography

Date GC-MS Run – date of performing the GC-MS run

GC Instrument – manufacturer and model of the GC instrument

GC Autosampler Model – manufacturer and model of the GC Autosampler

GC Column model – manufacturer and model of the column

GC Column type – column type e.g. normal-phase or reverse phase

Guard Column – manufacturer and model of the guard column

#Mass spectrometry

MS Scan polarity – choose between positive, negative, alternating, negative, NA or select Other and enter it

MS Scan mz range –mass to charge ratio scan range

MS Instrument – manufacturer and model of the MS instrument

MS Ion source – choose between electron ionization (EI), chemical ionization (CI), electrospray ionization (ESI), atmospheric pressure chemical ionization (APCI), atmospheric pressure photoionization (APPI), inductively coupled plasma ionization (ICP), matrix assisted laser desorption and ionization (MALDI) or select Other and enter it

Mass analyzer – choose between quadrupole mass filter, quadrupole ion trap, Fourier-transform ion cyclotron resonance (FT-ICR), orbitrap, time-of-flight (TOF) or select Other and enter it

Raw Spectral Data Files – link to files or directory with raw spectral data

#Other

Operator – name or acronym of the person doing the wet lab work
